# Amyloid-beta biomarkers in Braak stages and their predictive relationships with cognitive impairment: Support vector machine and deep learning approaches

**DOI:** 10.1101/2022.09.25.509432

**Authors:** Ahmed Taha, Bhavin Soni, Deepa S. Thakuri, Erin Ritter, Puskar Bhattarai, Ganesh B. Chand

## Abstract

Amyloid-beta (*Aβ*) and tau tangles are hallmarks of Alzheimer’s disease. *Aβ* distributions in the tau-defined Braak staging regions and their multivariate predictive relationships with mild cognitive impairment (MCI) are not known. In this study, we used PiB PET data from 60 participants (33 with MCI and 27 healthy controls (HC)), quantified *Aβ* as distribution volume ratio (DVR) in Braak regions, and compared between MCI and controls to test the hypothesis that DVR alters with declining cognition. We found elevated DVR in participants with MCI, especially in the spatial distribution of Braak stages III-IV and V-VII, while an alteration in Braak stage I-II was near the statistical significance. DVR markers correlated with cognitive status, especially in Braak stages III-IV and VI-V. To evaluate whether these markers are predictive of cognitive dysfunction, we designed support vector machine and artificial neural network models. These methods showed predictive multivariate relationships between *Aβ* makers of Braak regions and cognitive impairment. Overall, these results highlight the importance of computer-aided research efforts for understanding AD pathophysiology.

## 1. Introduction

Alzheimer’s disease (AD) is the most common type of dementia worldwide [1]. Rather than being a singular diagnosis, AD is a disease spectrum that begins with an asymptomatic phase or preclinical AD and extends to the severe symptomatic AD dementia [2, 3]. On that spectrum, mild cognitive impairment (MCI) represents the transitional state between normal cognitive changes of aging and early AD [4]. Amyloid-beta (*Aβ*) and tau have emerged as two important hallmarks for AD research, however their inter-relationships with MCI is not well understood. Specifically, regional *Aβ* markers based on the tau-defined topological regions and their multivariate predictive relationships with cognitive impairment is unclear yet.

In AD pathophysiology, the amyloid cascade hypothesis is the most studied one [5]. It postulates that the accumulation of the extracellular *Aβ* plaques is the primary event that leads to a sequence of intracellular neurofibrillary tangles accumulation, synaptic dysfunction, and gliosis; collectively, these events lead to symptomatic AD dementia in late stages of the disease [6]. Although further research challenged the causality of *Aβ* accumulation as a sole cause of such a sequence, some suggest that *Aβ* accumulation could be the vital ignition key to a more complicated patho-physiological process [7]. For decades, AD diagnosis has relied on the gold standard of postmortem histopathological diagnosis [8]. This approach has dramatically changed with the utilization of positron emission tomography (PET) imaging and the introduction of Pittsburgh Compound B (PiB); the first in-vivo *Aβ* radiotracer [9, 10]. The notion that *Aβ* accumulation starts decades before developing any symptoms with sparked the search for novel imaging and fluid biomarkers [11], including markers for tau neurofibrillary tangles, *Aβ* plaques, neurodegeneration, gliosis, and even for other comorbidities that frequently occur with AD [12]. Braak staging, introduced in 1991, is a method of classifying the degree of tau pathology in AD. Briefly, there are six stages; namely: I&II where the neurofibrillary tangles are limited to the entorhinal region, stages III & IV where tangles progress to include the para-hippocampus and other parts of the limbic regions, and finally stages V & VI where the plaques extend to include a much wider area of the neocortex [13–15].

The association between regional tau neurofibrillary tangles accumulation and cognitive decline is somewhat established [16–18]. In fact, some literature suggests the superiority of tau accumulation and tau-PET over *Aβ* accumulation and (*Aβ*)-PET as a predictor of cognitive deterioration [19, 20]. Moreover, to the best of our knowledge, the accumulation of *Aβ* plaques in the topographic map of Braak staging in persons with MCI is not well established. The temporal relationship between tau and *Aβ* accumulation is a matter of debate; however, in a cross-sectional study, Jack *et al.* [21] suggested that the transformation from being *Aβ* +ve and tau-ve to *Aβ* +ve and tau +ve is associated with the beginning of severe cognitive dysfunction. Nevertheless, the association between global and regional tau deposition topography and that of *Aβ* is yet to be discovered [22]. One of the critical challenges along that line is the extensive off-target binding of tau radiotracers[23, 24]. Thus, understanding *Aβ* distribution in the tau-pathology defined regions is an active research approach, and advanced machine learning algorithms might aid along that line. Specifically, whether regional *Aβ* measured in tau-neuropathology defined Braak stages have multivariate relationships with MCI and whether such relationships can be predicted (to some extent) using advanced machine learning algorithms remain to be investigated.

In this study, we examine the accumulation of *Aβ* in the regions where tau neurofibrillary tangles are expected to accumulate and its relationship with cognitive dysfunction. We hypothesize that *Aβ* accumulation in the different Braak stages differs in MCI compared to controls. We further hypothesize that the different Braak stages will have different associations with cognitive performance. To evaluate our hypothesis whether these regional metrics have multivariate associations with cognitive dysfunction and whether such associations is predictive in nature using quantitative modeling, we designed machine learning models and estimated cognitive impairment in MCI.

## 2. MATERIAL AND METHODS

### 2.1. Participants

Participants of this study were chosen from the Open Access Series of Imaging Studies (OASIS) dataset [25, 26] (https://www.oasis-brains.org). We used the data of 60 participants; 33 of them are people with MCI and 27 controls, with a mean age (SD) of 77.3 (6.68) in MCI and a slightly lower mean of 73.3 (7.48) in controls. 45.45% and 55.56% of participants were females in MCI an in controls respectively.Average MMSE (standard deviation) was 26.70 (2.64) in subjects with MCI and 28.70 (1.64) in controls. Data were acquired from the OASIS as per our previous work [27]. Study participants were given an MCI diagnosis based on the widely used clinical dementia rating (CDR) scale [28, 29]. CDR is utilized to assess cognitive and functional abilities in AD and other dementias; participants are given rates from 1-5 in six different domains: memory, orientation, problem solving and judgment, personal care, home and hobbies and function in community affairs. Data are collected from both the participants and a collateral source of information (such as a family member) all through a semi-structured interview. A global CDR of 0 means no dementia whereas as score of 0.5 – 1, 2, 3 indicated very mild, mild and sever dementia, respectively. In our study, MCI participants had a CDR score of 0.5-1. We only included participants with AD being their primary cause of dementia [28, 29]. In this study, we employed the Mini-Mental State Examination (MMSE) to assess participants’ cognition for illness severity metrics. MMSE assesses five areas of cognition: orientation, registration, attention and calculation, recall, and language [30]. The maximum score is 30; scoring 23 or less indicates cognitive impairment (Folstein et al., 1975). Having low scores on MMSE and reduced MMSE domains are associated with increased mortality risk [31].

### 2.2. Data acquisition and processing

Data acquisition and processing were performed as per our previous work [27]. In brief, bolus injection of [11]C-PiB, was followed by Dynamic PET 3D scans that were acquired for more than an hour, using ECATHR plus 962 PET scanner or Biograph 40 PET/CT scanner. All participants got T1-weighted brain MRI using 3T Biograph MR or 1.5T Vision, Siemens 3T TrioTim. Multi-atlas region Segmentation utilizing Ensembles of registration algorithms and parameters and locally optimal atlas selection (MUSE) was used for individual T1-MRI segmentation [27, 32, 33]. We quantified dynamic PET scans as the Distribution Volume Ratio (DVR) outcome [34], the most suitable one compared to other statistic approaches. One challenge to the appropriate quantification of PET images is the partial volume effect [35] which was accounted for in our study by using a state-of-the-art parallel level set (PLS) regularization-based partial volume correction method [36]. To harmonize multi-scanner datasets, we relied on our previously used and validated approach [27, 32] in which we used data from all study participants to compute the coefficients accounting for the effect of factors such as site, age and sex and apply them to each individual participant [37]. After correcting for the site, age, and sex covariates, we produced the harmonized DVR and used that in our analyses. Braak staging regions are showed in **Table** I.

**TABLE I:**
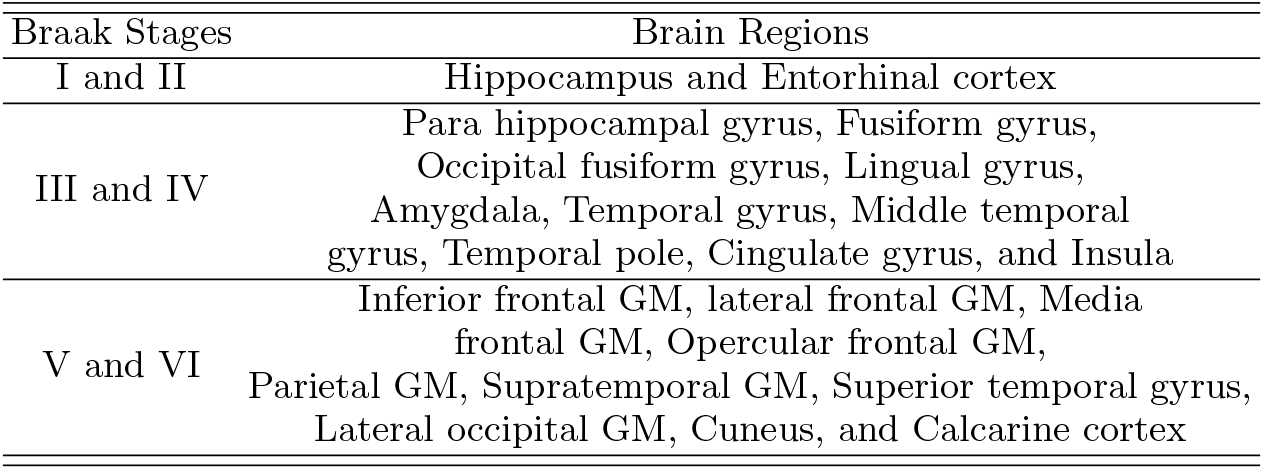
Brain regions of different Braak stages used in our study (GM: grey matter; brain regions represent both left and right hemispheres).

### 2.3. Machine learning based predictive modeling algorithms

To test if the *Aβ* markers (DVRs) in different Braak staging regions can help predict cognitive abilities (assessed via MMSE) in participants with MCI, the DVRs were used as the input variables, and the MMSE scores as output (response) variable. For our study, we employed two machine learning algorithms, especially support vector machine (SVM) consistent with our previous studies [27, 38] and artificial neural network (ANN) to evaluate the predictions of MMSE. For the SVM model (Fig. 1), we used 10-fold cross validation (CV). The SVM model was designed using the sci-kit learn package in python 3.0.1 [39]. For this approach, the dataset was divided into 10 equal folds. The model uses 9 folds as training set, and the remaining fold as the test set. This process is repeated for all the possible combinations of the 10-fold data. For this model, the radial bias function (RBF) kernel was used [40], and model parameters were optimized for each training set using grid search CV. The same SVM model designs were employed in each of the Braak stages and their combinations for the comparison while predicting MMSE. For the ANN model (Fig. 2), we used a 10-fold CV approach consistent with above. This model was implemented on Python 3.0.1 using TensorFlow [41] and Keras [42]. In our ANN model, we had one input layer, three hidden layers, and one output layer. Standard scalar was used before model training. For our input layer, 40 units were used, along with the rectified linear unit (relu) activation layer, and batch normalization at the end. For our first, second and third hidden layers, 30, 20 and 10 units were used, respectively. The sigmoid activation function was used in each hidden layer and batch normalization was performed after each hidden layer. Dropouts of 0.2 were used after the first and the second hidden layer to ensure there is no overfitting in the modeling analyses. For our output function, we used sigmoid activation function. This ANN model was implemented on the four Braak Staging Combinations like SVM.

**FIG. 1:**
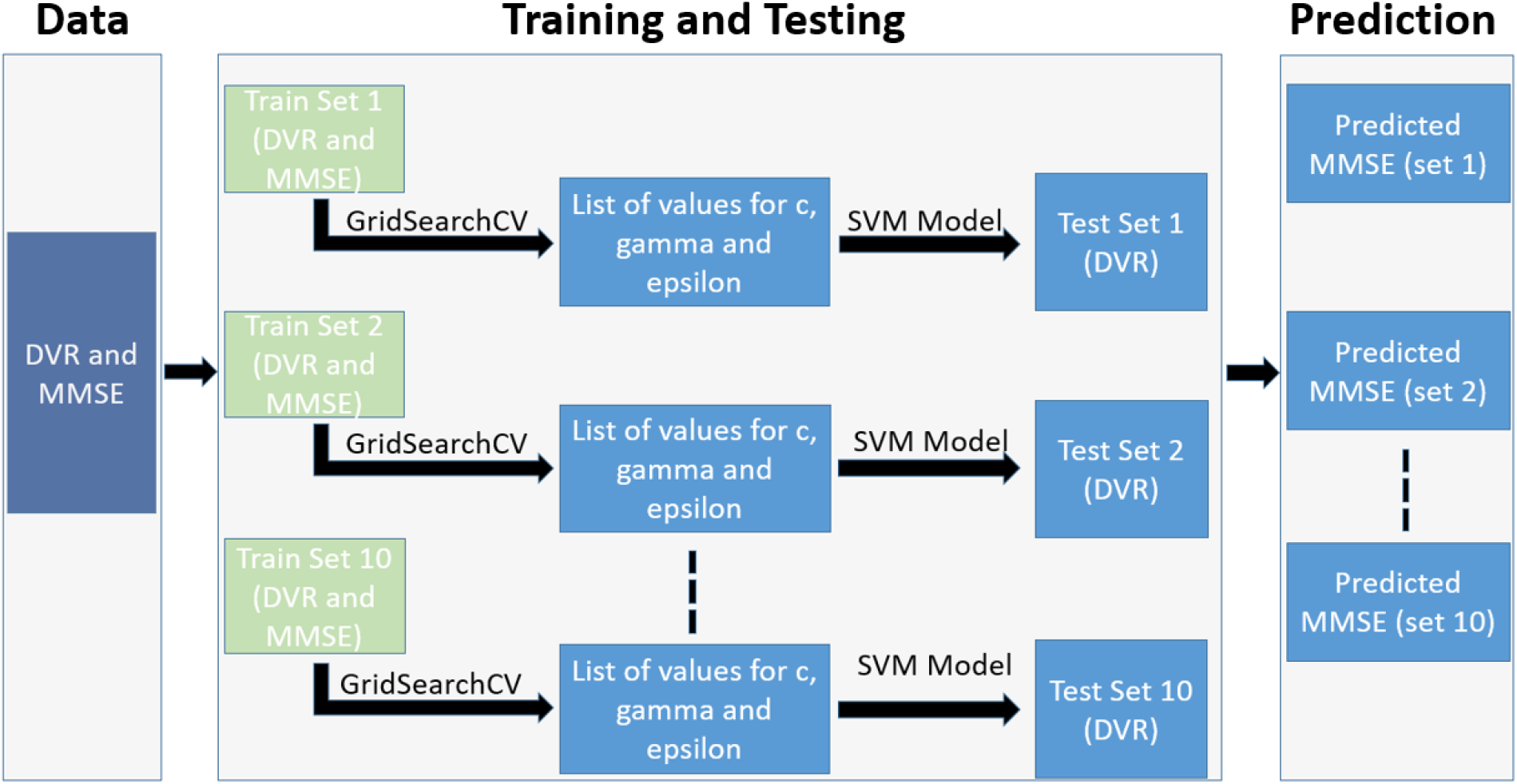
An illustration for the Support Vector Machine (SVM) modeling approach for predicting MMSE using DVRs as the feature variables. DVRs and MMSE of MCI from each of the Braak stages were fed into the SVM model. Grid Search CV was used to perform 10-fold cross validation on the entire data set; dividing our data into train set and test set in 10 folds. For each of our combinations, we used a list of c, epsilon, and gamma values, which were optimized and the optimal values for these hyperparameters were used on the testing datasets. During testing, only the DVR’s were fed to the SVM model.

**FIG. 2:**
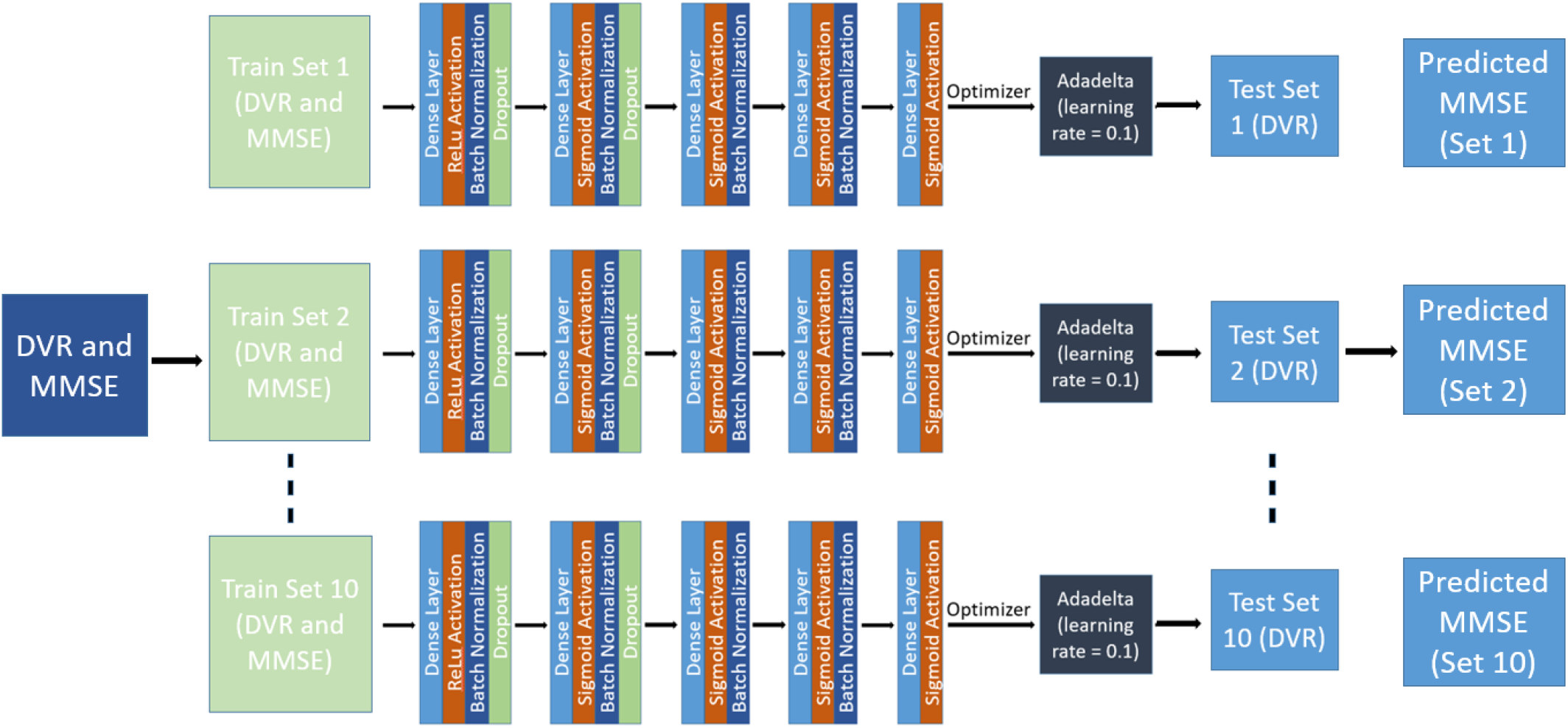
An illustration for the Artificial Neural Network (ANN) modeling approach for predicting MMSE using DVRs as feature variables. For the input layer of our ANN approach, we used 40 units, followed by relu activation and batch normalization, and a dropout layer of 0.2. For our first hidden layer, we used 30 units, and a relu activation, followed by relu activation layer, and a dropout of 0.2. For our second hidden layer, we used 20 units, followed by a sigmoid activation, and batch normalization. Our last hidden layer had 10 units, a sigmoid activation layer, followed by batch normalization. For our output layer, we used sigmoid activation.

### 2.4. Statistical analysis

To differentiate between the DVR medians in various Braak stages, in both controls and MCI patients, we used the Mann Whitney U test [43]. Spearman correlation was used to examine the relationship between MMSE and DVR. Spearman correlation was also used to determine the predictive associations between actual MMSE and predicted MMSE using machine learning algorithms. To adjust the statistical significance for multiple comparisons, a false discovery rate (FDR) [44] approach was implemented.

## 3. RESULTS

### 3.1. Comparison of DVRs between MCI and controls

We compared the medians of DVRs (composite of all regions within each Braak stage) between 33 participants with MCI and 27 controls (Fig. 3). We found highly elevated DVR in MCI participants, particularly in the regions of spatial distribution of Braak stages III-IV and V-VI. Interestingly, in stages I-II,the median DVR for controls was somewhat higher (1.009) compared to that of people with MCI 0.935, but the difference was minimal. In stages III-IV, the median DVR for MCI (1.004) is higher than for controls (0.905, *FDR* – *p* = 0.008) and for stages V-VI(*FDR* – *p* = 0.002) they are 1.058 and 0.903 respectively. Combining all stages, we found that in general, people with MCI have a marginally higher statistically significant (*FDR* – *p* = 0.002) DVR than that of controls, with medians of 0.972 and 0.935 respectively.

**FIG. 3:**
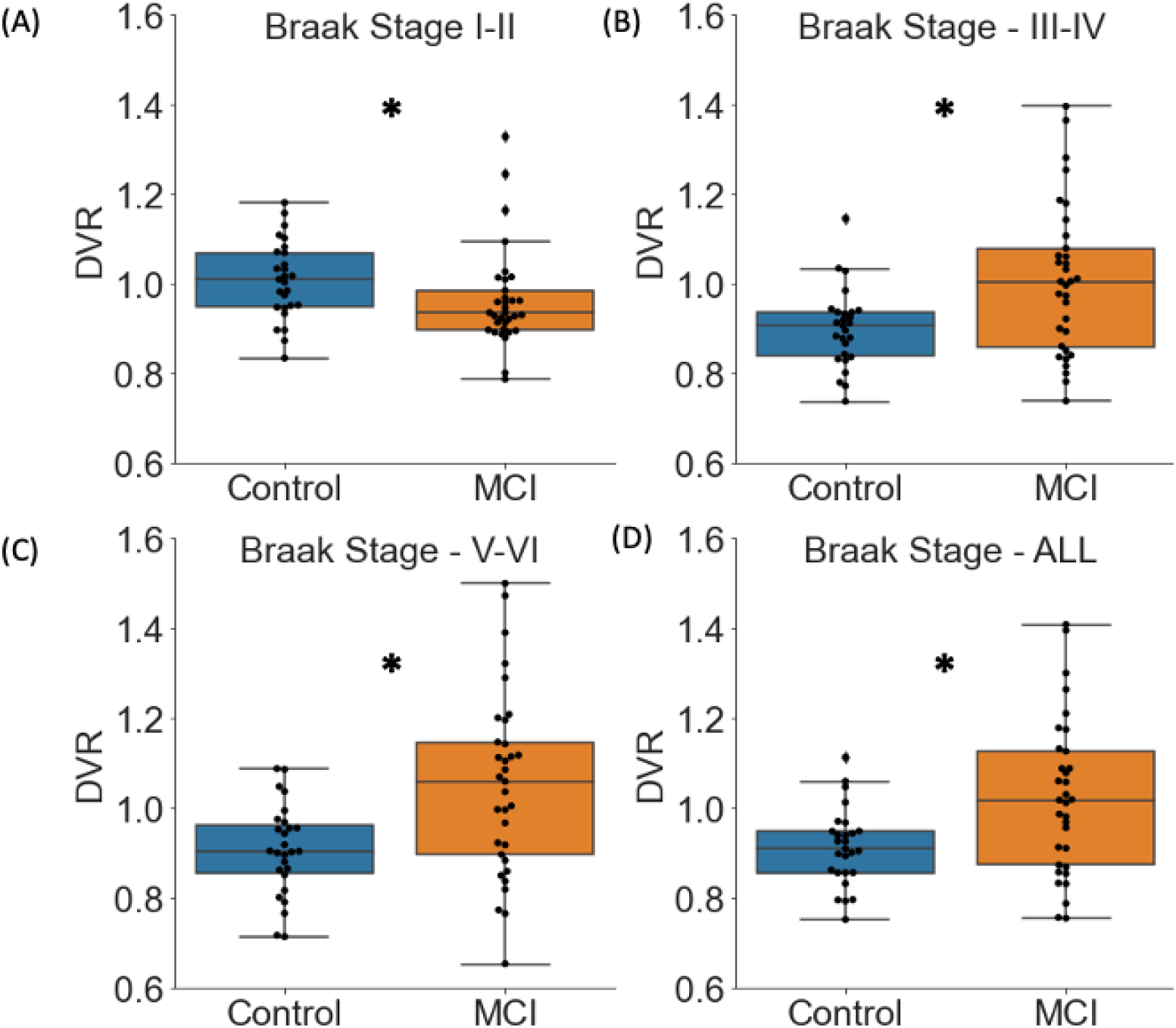
Boxplots and scatterplots representing Distribution Volume Ratio (DVRs) of PET *Aβ* scans in participants with mild cognitive impairment (MCI) and controls. (A-C). Median 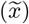, (\**FDR* corrected p-value). Braak Staging I-II, controls 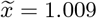, MCI 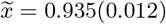. Braak staging III-iV, controls 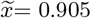, MCI 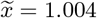, (0.008). Braak Staging V and VI controls 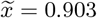, MCI 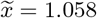.(0.002). (D) represents DVR for all the brain regions associated with all the six Braak stages. controls 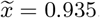, MCI 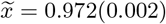.

### 3.2. Associations between DVR and MMSE

Spearman’s correlation was calculated between DVR and MMSE in MCI and controls participants together (Fig. 4). Correlation strength was very weak in early stages I-II (*ρ* = 0.063; *FDR* – *p* = 0.698). Correlation strength was somewhat stronger and highly significant in stage III-IV (*ρ* = –0.423; FDR-p=0.003), stage V-VI (*ρ* = –0.440; *FDR* ‒ *p* = 0.003), and all stages together (*ρ* = –0.429; *FDR* – *p* = 0.003). We further studied the associations between DVR and MMSE only in MCI. In the data of 33 participants with MCI (Fig. 5), the associations between DVR and MMSE were especially stronger in later stages: stage I-II (*ρ* = 0.017; FDR – p = 0.924), stage III-IV (*ρ* = –0.451; FDR – *p* = 0.011), stage V-VI (*ρ* = –0.474; *FDR* – *p* = 0.012), and all stages (*ρ* = –0.496; *FDR* – *p* = 0.005).

**FIG. 4:**
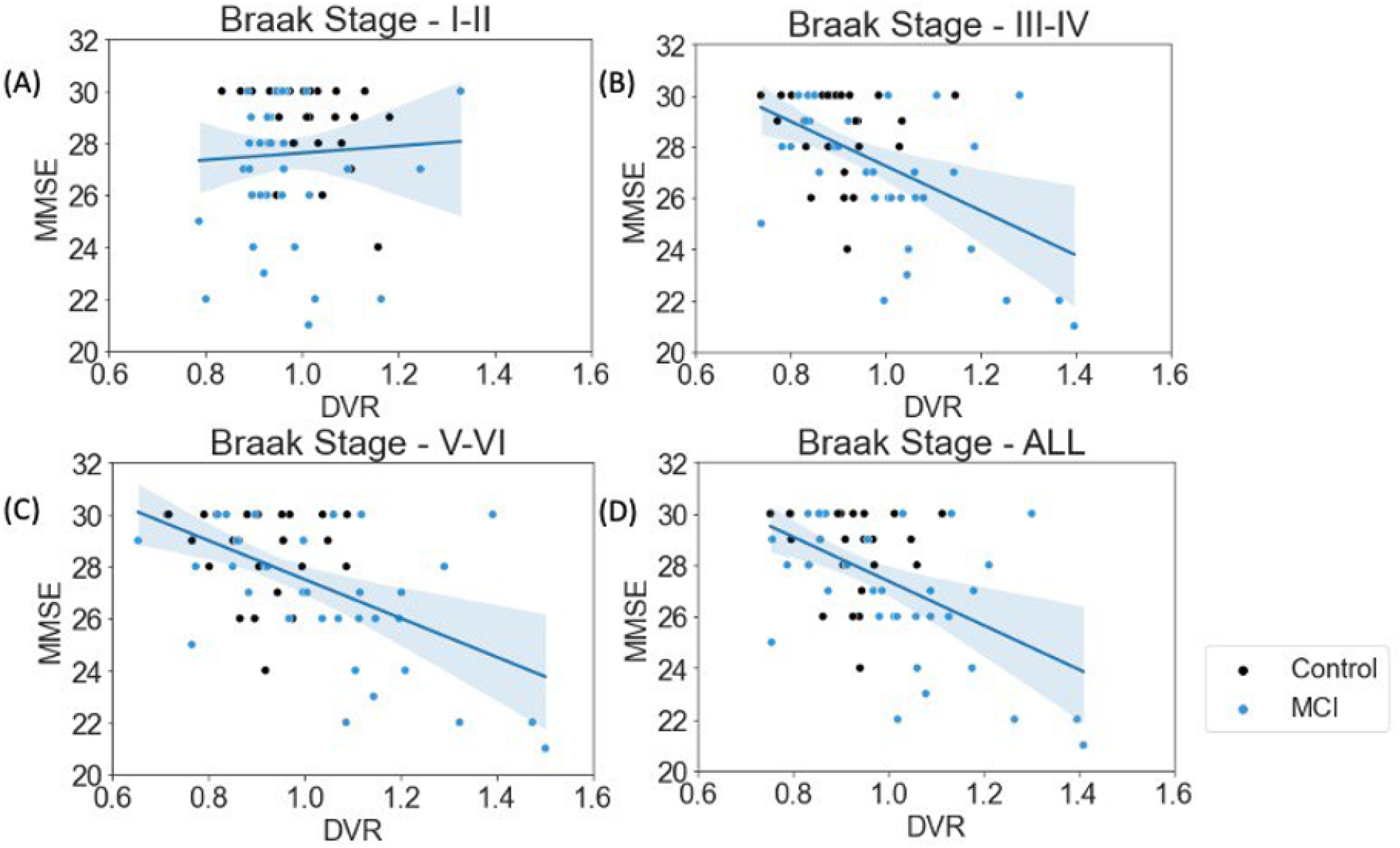
Correlation plots between the distribution volume ratio (DVR) of different Braak Stage Regions and the Mini-Mental State Examination (MMSE) Score in all the participants. (*FDR corrected p-value) (A) denotes that there is a weak positive correlation between the DVRs of regions associated with Braak Staging I and II, while (B-D) denote that there is a negative correlation between the DVRs of regions related with Braak Stage III-IV, V-VI, and when all regions are considered (* : *FDR* – *p* < 0.05). The shadowed area in the plots (A-D) denote the confidence interval. Stages I-II, *ρ* = 0.063(0.698). III-IV, *ρ* = –0.423(0.003). V-VI, *ρ* = –0.440(0.003). In all Braak stages, *ρ* = –0.429(0.003).

**FIG. 5:**
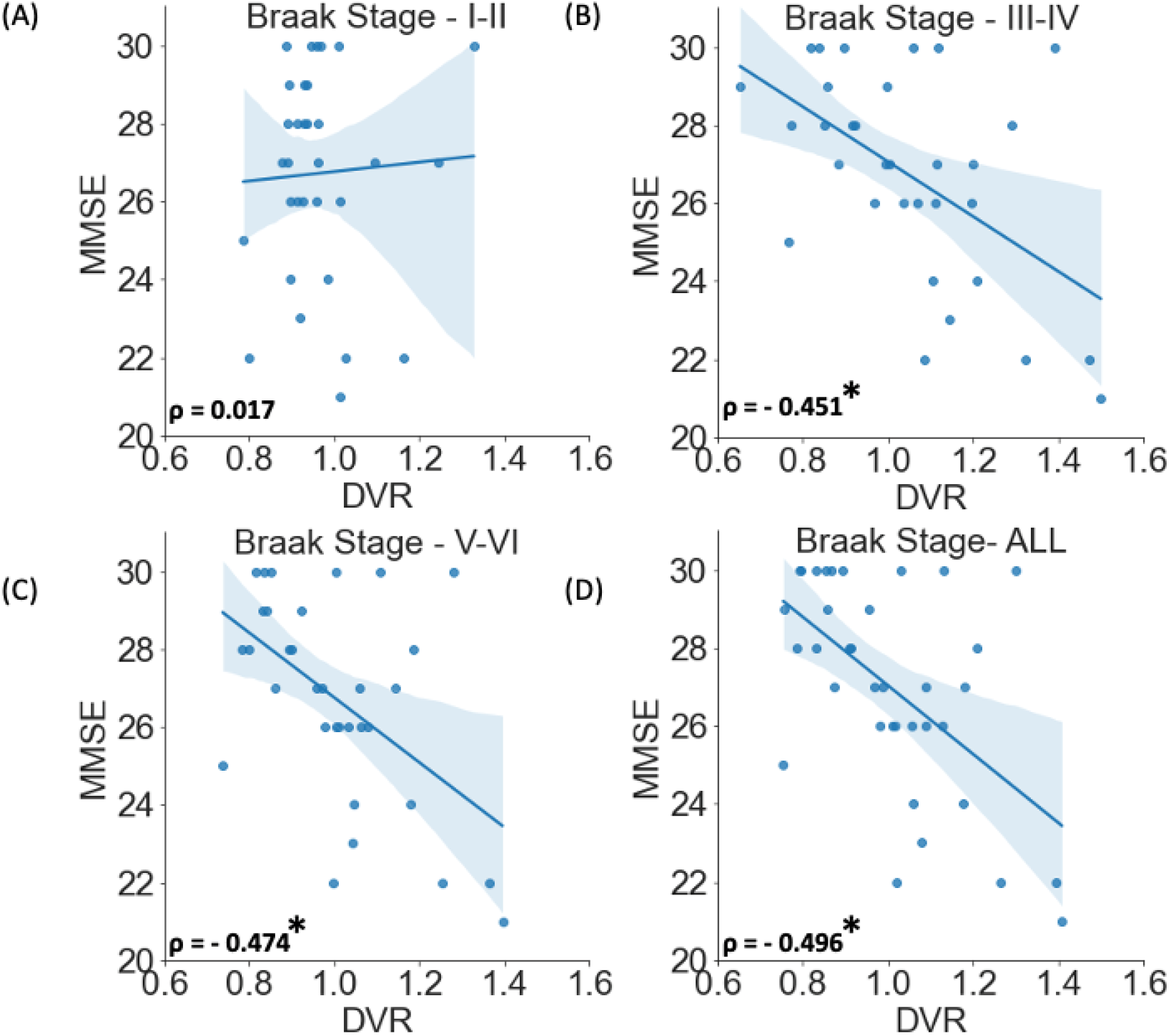
Correlation plots between the distribution volume ratio (DVR) of different Braak Stage Regions and the Mini-Mental State Examination (MMSE) score in only the MCI Participants. (A) denotes that there is a weak positive correlation between the DVRs of regions associated with Braak Staging I and II, while (B-D) denote that there is a negative correlation between the DVRs of regions related with Braak Stage III-IV, V-VI, and when all regions are taken into account (* : *FDR* – *p* < 0.05). The shadowed area in the plots (A-D) denote the confidence interval. Stages I-II, *ρ* = 0.017(0.924), III-IV, with *ρ* = –0.451(0.011), V-VI; *ρ* = –0.474(0.012). In all stages combined, correlation was *ρ* = –0.496, (0.005).

## 4. Machine learning based MMSE predictions

SVM and ANN models were used to evaluate if the regional DVR values can predict MMSE scores in participants with MCI in different Braak Stages, and to assess which model is more accurate in prediction of the same. For our SVM models using the DVRs for regions in Braak Staging I and II, we found out that the predicted MMSE and actual MMSE have a very low correlation in patients with MCI (*ρ*= 0.069; *p* > 0.05) (Fig. 6). However, for Braak Staging III and IV, using DVRs as predictors for MMSE provided with a high correlation between predicted and actual MMSE (*ρ* = 0.530; FDR-p =0.004). Similarly, for DVRs of Braak Staging V and VI, the correlation between actual and predicted MMSE was high (*ρ* = 0.371; *FDR* – *p* = 0.041). The DVRs for all the Braak staging regions were also used to check for using SVM in predicting MMSE. The correlation for predicted MMSE vs. actual MMSE for this case was high (*ρ* = .538; *FDR* – *p* = 0.004). The ANN models performed generally better than the SVM models in using DVRs to predict the MMSE in participants with MCI (Fig. 7). Braak staging I and II (*ρ* = 0.369; *FDR* – *p* = 0.041), Braak Staging III and IV (*ρ* = 0.594; FDR-p =0.003), Braak Staging V and VI (*ρ* = 0.543; *FDR* – *p* = 0.004), and all Braak staging combined (*ρ* = 0.665; *FDR* – *p* = 0.001) had statistically significant correlation between predicted MMSE and actual MMSE.

**FIG. 6:**
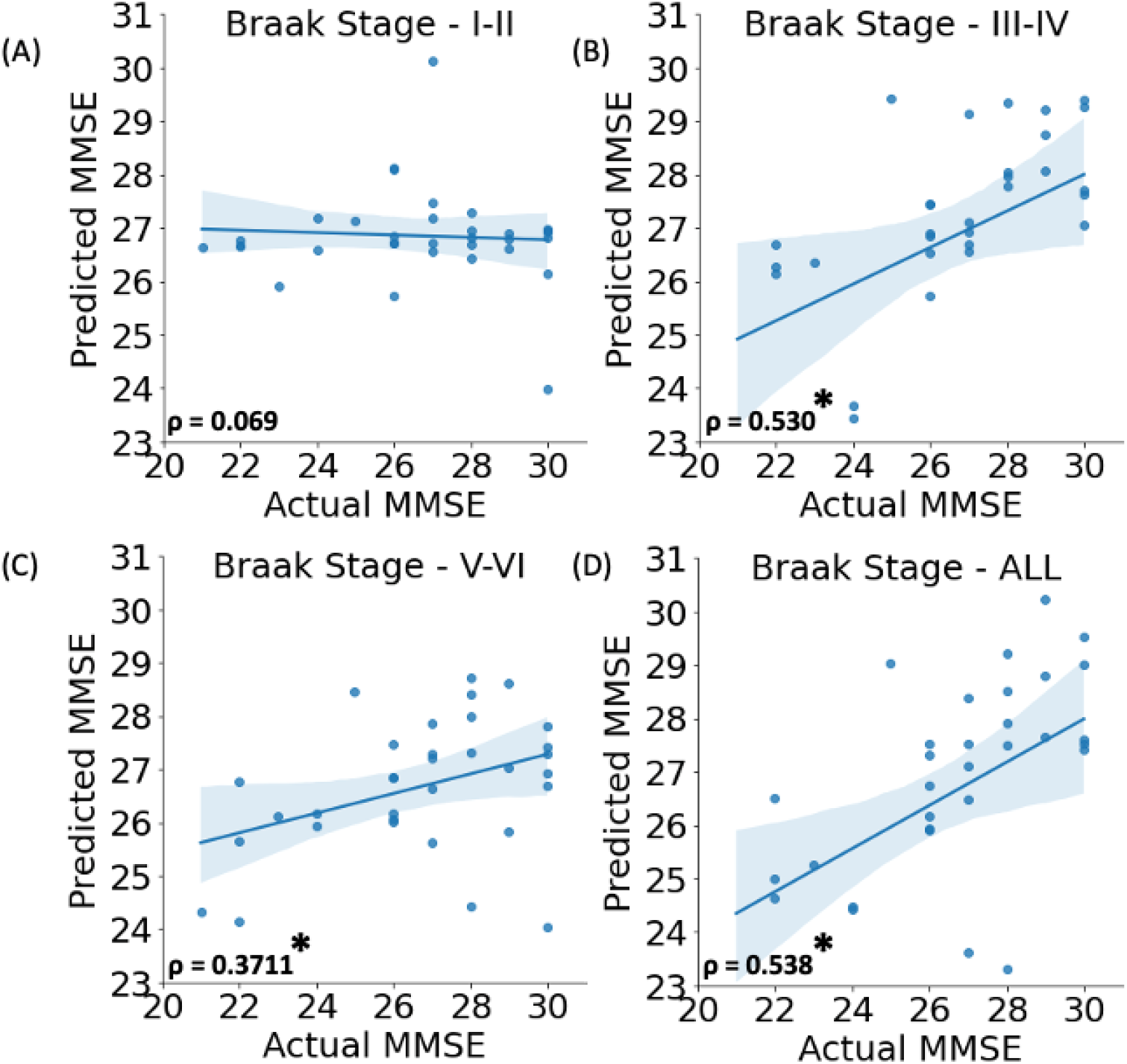
Correlation plots between the predicted Mini-Mental State Examination (MMSE) score and the actual MMSE score in participants with mild cognitive impairment (MCI) using Support Vector Machines (SVM). All the measurements have a positive correlation, but only (B-D) have a statistical significance (* : *FDR* – *p* < 0.05) between predicted MMSE and actual MMSE scores. The correlation was somewhat higher when all the features are considered to estimate MMSE (Fig. D). The shadowed area in the plots (A-D) denote the confidence interval. I-II, (*ρ*= 0.069; *p* > 0.05). III and IV, (*ρ* = 0.530; *p* < 0.05). Similarly, for DVRs of V-VI, (*ρ* = 0.3711; *p* < 0.05). All the Braak staging regions (*ρ*= .538; *p* < 0.05).

**FIG. 7:**
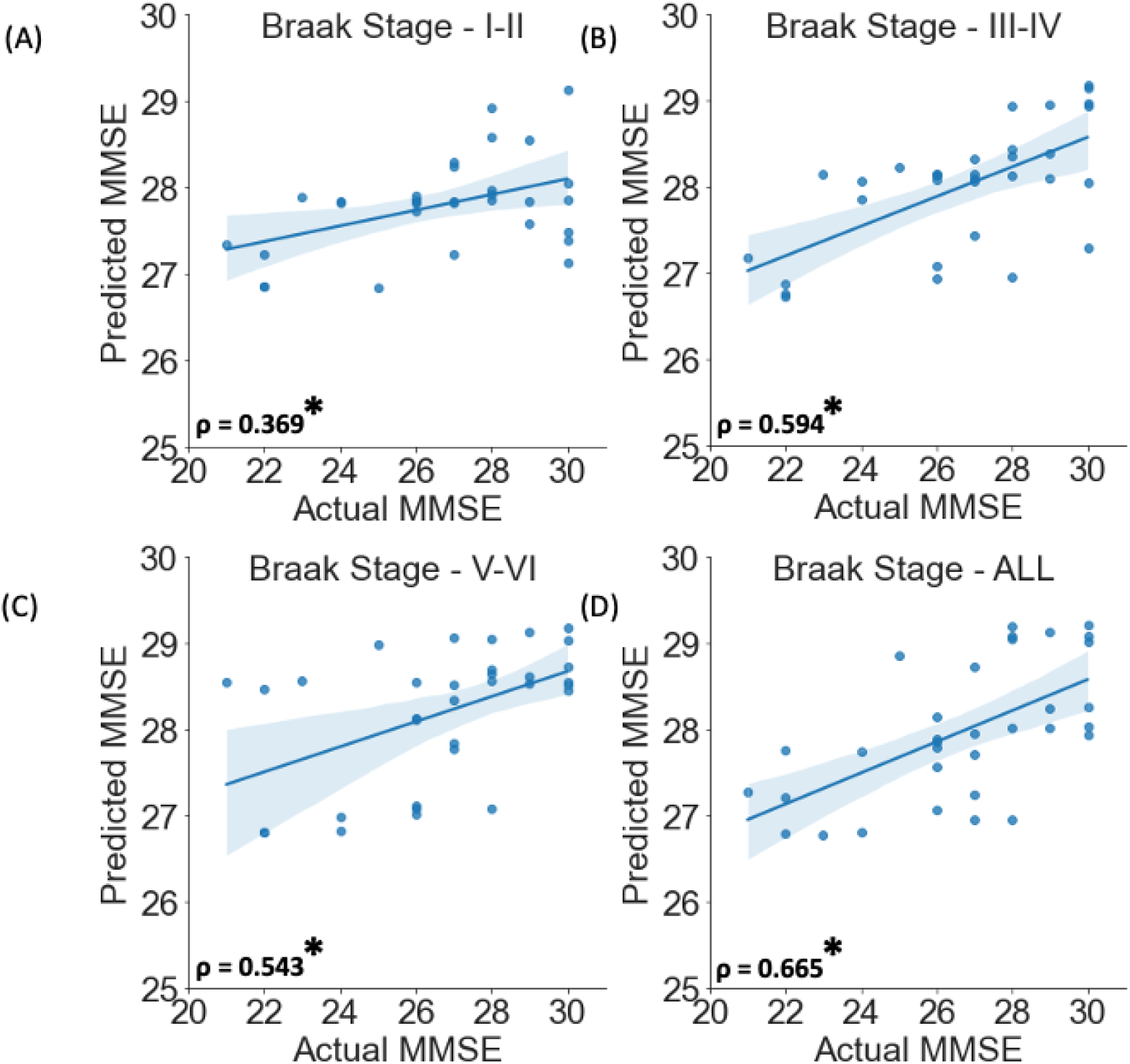
Correlation plots between the predicted Mini-Mental State Examination (MMSE) score and the actual MMSE score in participants with mild cognitive impairment (MCI) using Artificial Neural Networks (ANN). All the measurements have a positive correlation, but only (B-D) have a statistical significance (* : *FDR* – *p* < 0.05) between predicted and actual MMSE scores. The correlation is higher when all the features were considered to predict MMSE (Fig. D). The shadowed area in the plots (A-D) denote the confidence interval. Stages I-II (*ρ* = 0.369; *p* < 0.05), III-IV (*ρ* = 0.594; *p* < 0.05), Braak Staging V-VI (*ρ* = 0.543; *p* < 0.05) and all staging areas combined (*ρ* = 0.665; *p* < 0.05).

## 5. DISCUSSION

In this study, we quantified [11]C-PiB as DVRs in the spatial distribution of Braak stages and compared DVRs between participants with MCI and controls. Our results showed higher DVRs in persons with MCI compared to controls, especially in the latter stages. Furthermore, in participants with MCI, we found a significant correlation between DVRs and MMSE scores particularly in stages III-IV. Moreover, AAN and SVM predictive modeling approaches successfully predicted MMSE in people with MCI, especially starting at stages III-IV, and demonstrated that there are multivariate predictive relationships of regional accumulation of *Aβ* in the spatial distribution of Braak staging regions and cognitive dysfunction.

Previous literature suggests that the relationship between *Aβ* and tau accumulation is not fully understood; however, there is some consensus that *Aβ* is an upstream trigger of tau accumulation, and in-vivo studies have shown that *Aβ* potentiates the toxicity of tau [45–47]. Our approach of measuring *Aβ* in the spatial distribution of tau, marked by Braak staging boundaries, adds to the efforts of defining biomarkers to early detect cognitive decline, without facing off-target challenges associated with tau-PET, and hopefully aid in developing computer-assisted diagnostic and prognostic approaches. Our reasoning is also different from what has been established as a distinct spatial accumulation pattern for *Aβ* [10, 47]. In our study, the clear difference between DVRs in controls and MCI participants in different stages supports the hypothesis that regional *Aβ* accumulation is a useful and sensitive marker for disease progression, even more useful than the other biomarkers [48]. The effect of neuropathological changes on cognitive decline in MCI is complex, and the transition from normal cognition to MCI is complicated by the effects of age, apolipoprotein E e4 gene carriership [49], and protective factors such as the cognitive reserve [50] with age being a more significant factor than neurofibrillary pathological changes [51]. However, controlling for age had a smaller effect on cognitive deterioration when the accumulation of amyloid was measured [52]. Interestingly, it has been shown that amyloid is a sensitive predictive factor for tau accumulation [53]. Furthermore, Welikovitch et al. [54] proposed that accumulation of soluble forms of *Aβ* precedes the accumulation of insoluble forms, this accumulation was not related to any cognitive impairment though [54]. In contrast, it is well-established that the hippocampus and entorhinal cortex, which represent the topographic areas of Braak stages I-II, are the first to show significant changes with cognitive decline, in the form of neurofibrillary tangles [13, 55, 56], and in the form of atrophic changes, with more atrophy in the entorhinal cortex compared to the hippocampus [57]. Hippocampus and its impaired functional roles with other brain regions are widely reported in MCI [58]. Other brain regions that are part of Braak stages have also shown broad functional relationships with cognition in health and MCI [59–64].

Furthermore, the significant correlation between DVRs, especially in late stages (III-VI), and MMSE in people with MCI further supports the hypothesis that more *Aβ* accumulation leads to worse cognitive abilities, which eventually leads to higher risk of developing AD dementia. The correlation might also help clarifying the relationships between *Aβ* accumulation and tau, and their mutual effect on cognitive abilities and disease progression in later stages of the disease, i.e., symptomatic AD. It was noteworthy to find out that the correlation between DVRs and MMSE scores was not as strong in the spatial distribution of the early Braak stages, namely I-II. It was also quite surprising to see a slight higher DVRs of controls than MCI, although small effect size. One of the possible reasons is that some of controls are in almost at the borderline to meet the clinical diagnosis of MCI and/or about transitioning to MCI phase. Using ANN, and SVM models, we were able to reliably predict cognitive impairment in MCI population starting at Braak staging III-IV. Even with a rather limited data and a small study sample; the success of these models in predicting the MMSE scores based on the DVR only is a highly encouraging proof-of-concept approach. The results point to the significance machine learning algorithms to help understand and diagnose both MCI and AD, this conclusion agrees with previously published work on the reliability of machine learning models [25, 65, 66].

Some Limitations to this study need to be mentioned. The relatively small number of both participants with MCI and controls limits our approach, however it is a motive to nclude more participants in future studies to replicate our findings. Furthermore, the concept of *Aβ* accumulation needs to be tested on other brain regions since we only used the distribution of tau neurofibrilly tangles. Nonetheless, we think that the predictions can be improved in a bigger sample. Future work will also perform a head-to-head comparison between tau and *Aβ* accumulations in the same brain regions. Finally, we only applied our machine learning algorithms in the Braak staging regions; however, we suspect that combining other regions would improve the ability of our machine learning approaches to predict MMSE successfully in participants with MCI and probably in other populations.

In conclusions, the observed difference of *Aβ* accumulation in participants with MCI versus controls, especially in later stages of the spatial distribution of Braak stages, adds to the understanding of the temporal relationship between tau and *Aβ* accumulation. Moreover, the association between *Aβ* DVR and cognitive dysfunction (measured by MMSE) is indicative of the importance of *Aβ* as a biomarker for MCI. Finally, the prediction of cognitive dysfunction based on the DVRs of participants with MCI strengthens the role of the advanced machine learning modeling algorithms in helping computer-aided diagnose and predict the prognosis of neurodegenerative disorders.

## Acknowledgments

Mallinckrodt Institute of Radiology funding supports the senior author GBC.

## References

[1] J. Gaugler, B. James, T. Johnson, J. Reimer, M. Solis, J. Weuve, R. F. Buckley, and T. J. Hohman, Alzheimers & Dementia 18, 700 (2022).

[2] Y. Deschaintre, F. Richard, D. Leys, and F. Pasquier, Neurology 73, 674 (2009).

[3] A. S. Fleisher, K. Chen, Y. T. Quiroz, L. J. Jakimovich, M. G. Gomez, C. M. Langois, J. B. Langbaum, A. Roontiva, P. Thiyyagura, W. Lee, et al., JAMA neurology 72, 316 (2015).

[4] R. C. Petersen, R. Doody, A. Kurz, R. C. Mohs, J. C. Morris, P. V. Rabins, K. Ritchie, M. Rossor, L. Thal, and B. Winblad, Archives of neurology 58, 1985 (2001).

[5] C. Reitz, International journal of Alzheimer’s disease 2012 (2012).

[6] J. A. Hardy and G. A. Higgins, Science 256, 184 (1992).

[7] E. S. Musiek and D. M. Holtzman, Nature neuroscience 18, 800 (2015).

[8] A. Alzheimer, R. A. Stelzmann, H. N. Schnitzlein, and F. R. Murtagh, Clinical anatomy 8, 429 (1995).

[9] A. D. Cohen and W. E. Klunk, Neurobiology of disease 72, 117 (2014).

[10] W. E. Klunk, H. Engler, A. Nordberg, Y. Wang, G. Blomqvist, D. P. Holt, M. Bergström, I. Savitcheva, G.-F. Huang, S. Estrada, et al., Annals of Neurology: Official Journal of the American Neurological Association and the Child Neurology Society 55, 306 (2004).

[11] K. Blennow and H. Zetterberg, Journal of internal medicine 284, 643 (2018).

[12] H. Zetterberg and B. B. Bendlin, Molecular psychiatry 26, 296 (2021).

[13] H. Braak and E. Braak, Acta neuropathologica 82, 239 (1991).

[14] M. Kim, S. J. Kim, J. E. Park, J. Yun, W. H. Shim, J. S. Oh, M. Oh, J. H. Roh, S. W. Seo, S. J. Oh, et al., Scientific reports 11, 1 (2021).

[15] M. Schöll, S. N. Lockhart, D. R. Schonhaut, J. P. O’Neil, M. Janabi, R. Ossenkoppele, S. L. Baker, J. W. Vogel, J. Faria, H. D. Schwimmer, et al., Neuron 89, 971 (2016).

[16] D. A. Bennett, J. A. Schneider, R. S. Wilson, J. L. Bienias, and S. E. Arnold, Archives of neurology 61, 378 (2004).

[17] R. Ossenkoppele, D. R. Schonhaut, M. Schöll, S. N. Lockhart, N. Ayakta, S. L. Baker, J. P. O’Neil, M. Janabi, A. Lazaris, A. Cantwell, J. Vogel, M. Santos, Z. A. Miller, B. M. Bettcher, K. A. Vossel, J. H. Kramer, M. L. Gorno-Tempini, B. L. Miller, W. J. Jagust, and G. D. Rabinovici, Brain 139, 1551 (2016).

[18] L. Wang, T. L. Benzinger, Y. Su, J. Christensen, K. Friedrichsen, P. Aldea, J. McConathy, N. J. Cairns, A. M. Fagan, J. C. Morris, et al., JAMA neurology 73, 1070 (2016).

[19] D. Biel, M. Brendel, A. Rubinski, K. Buerger, D. Janowitz, M. Dichgans, and N. Franzmeier, Alzheimer’s research & therapy 13, 1 (2021).

[20] P. Giannakopoulos, F. Herrmann, T. Bussière, C. Bouras, E. Kövari, D. Perl, J. Morrison, G. Gold, and P. Hof, Neurology 60, 1495 (2003).

[21] C. R. Jack Jr, H. J. Wiste, H. Botha, S. D. Weigand, T. M. Therneau, D. S. Knopman, J. Graff-Radford, D. T. Jones, T. J. Ferman, B. F. Boeve, et al., Brain 142, 3230 (2019).

[22] S. N. Lockhart, M. Schöll, S. L. Baker, N. Ayakta, K. N. Swinnerton, R. K. Bell, T. J. Mellinger, V. D. Shah, J. P. O’Neil, M. Janabi, et al., Neuroimage 150, 191 (2017).

[23] C. Groot, S. Villeneuve, R. Smith, O. Hansson, and R. Ossenkoppele, Journal of Nuclear Medicine 63, 20S (2022).

[24] R. Harada, N. Okamura, S. Furumoto, and K. Yanai, Frontiers in neuroscience 12, 585 (2018).

[25] P. J. LaMontagne, T. L. Benzinger, J. C. Morris, S. Keefe, R. Hornbeck, C. Xiong, E. Grant, J. Hassenstab, K. Moulder, A. G. Vlassenko, et al., MedRxiv (2019).

[26] D. S. Marcus, A. F. Fotenos, J. G. Csernansky, J. C. Morris, and R. L. Buckner, Journal of cognitive neuroscience 22, 2677 (2010).

[27] G. B. Chand, D. S. Thakuri, and B. Soni, Journal of Neuroimaging 32, 728 (2022).

[28] M. J. C., Neurology 43, 2412 (1993).

[29] J. C. Morris, M. Storandt, J. P. Miller, D. W. McKeel, J. L. Price, E. H. Rubin, and L. Berg, Archives of neurology 58, 397 (2001).

[30] C. D. Patnode, L. A. Perdue, R. C. Rossom, M. C. Rushkin, N. Redmond, R. G. Thomas, and J. S. Lin, Jama 323, 764 (2020).

[31] Y. Su, J. Dong, J. Sun, Y. Zhang, S. Ma, M. Li, A. Zhang, B. Cheng, S. Cai, Q. Bao, S. Wang, and P. Zhu, BMC geriatrics 21, 1 (2021).

[32] G. B. Chand, D. B. Dwyer, G. Erus, A. Sotiras, E. Varol, D. Srinivasan, J. Doshi, R. Pomponio, A. Pigoni, P. Dazzan, et al., Brain 143, 1027 (2020).

[33] J. Doshi, G. Erus, Y. Ou, S. M. Resnick, R. C. Gur, R. E. Gur, T. D. Satterthwaite, S. Furth, C. Davatzikos, A. N. Initiative, et al., Neuroimage 127, 186 (2016).

[34] J. Logan, N. D. Volkow, J. S. Fowler, G.-J. Wang, S. L. Dewey, R. MacGregor, D. Schlyer, S. J. Gatley, N. Pappas, P. King, et al., Journal of Cerebral Blood Flow & Metabolism 14, 995 (1994).

[35] W. W. Moses, Nuclear Instruments and Methods in Physics Research Section A: Accelerators, Spectrometers, Detectors and Associated Equipment 648, S236 (2011).

[36] Y. Zhu, M. Bilgel, Y. Gao, O. G. Rousset, S. M. Resnick, D. F. Wong, and A. Rahmim, Physics in Medicine & Biology 66, 145003 (2021).

[37] G. B. Chand, P. Singhal, D. B. Dwyer, J. Wen, G. Erus, J. Doshi, D. Srinivasan, E. Mamourian, E. Varol, A. Sotiras, G. Hwang, P. Dazzan, R. S. Kahn, H. G. Schnack, M. V. Zanetti, E. Meisenzahl, G. F. Busatto, B. Crespo-Facorro, C. Pantelis, S. J. Wood, C. Zhuo, R. T. Shinohara, H. Shou, Y. Fan, N. Koutsouleris, A. N. Kaczkurkin, T. M. Moore, A. Verma, M. E. Calkins, R. E. Gur, R. C. Gur, M. D. Ritchie, T. D. Satterthwaite, D. H. Wolf, and C. Davatzikos, American Journal of Psychiatry 179, 650 (2022).

[38] G. B. Chand, M. Habes, S. Dolui, J. A. Detre, D. A. Wolk, and C. Davatzikos, Journal of neuroscience methods 331, 108528 (2020).

[39] A. Abraham, F. Pedregosa, M. Eickenberg, P. Gervais, A. Mueller, J. Kossaifi, A. Gramfort, B. Thirion, and G. Varoquaux, Frontiers in Neuroinformatics 8 (2014).

[40] N. Cristianini, J. Shawe-Taylor, et al., (2000).

[41] M. Abadi, A. Agarwal, P. Barham, E. Brevdo, Z. Chen, C. Citro, G. S. Corrado, A. Davis, J. Dean, M. Devin, et al., arXiv preprint arXiv:1603.04467 (2016).

[42] F. Chollet, “keras,” (2015).

[43] M. P. Fay and M. A. Proschan, Statistics surveys 4, 1 (2010).

[44] Y. Benjamini and Y. Hochberg, Journal of the Royal statistical society: series B (Methodological) 57, 289 (1995).

[45] G. S. Bloom, JAMA neurology 71, 505 (2014).

[46] J. Lewis, D. W. Dickson, W.-L. Lin, L. Chisholm, A. Corral, G. Jones, and S.-H. Yen, Science 293, 1487 (2001).

[47] D. R. Thal, U. Rüb, M. Orantes, and H. Braak, Neurology 58, 1791 (2002).

[48] J. Pfeil, M. C. Hoenig, E. Doering, T. van Eimeren, A. Drzezga, G. N. Bischof, A. D. N. Initiative, et al., Neurobiology of aging 106, 119 (2021).

[49] B. Stephan, S. Hunter, D. Harris, D. Llewellyn, M. Siervo, F. Matthews, and C. Brayne, Molecular psychiatry 17, 1056 (2012).

[50] S. Mazzeo, S. Padiglioni, S. Bagnoli, L. Bracco, B. Nacmias, S. Sorbi, and V. Bessi, Journal of Neurology 266, 487 (2019).

[51] E. J. Mufson, M. Malek-Ahmadi, N. Snyder, J. Ausdemore, K. Chen, and S. E. Perez, Neurobiology of aging 43, 101 (2016).

[52] S. Villeneuve, B. R. Reed, M. Wirth, C. M. Haase, C. M. Madison, N. Ayakta, W. Mack, D. Mungas, H. C. Chui, C. DeCarli, et al., Neurology 82, 761 (2014).

[53] S. L. Leal, S. N. Lockhart, A. Maass, R. K. Bell, and W. J. Jagust, Journal of Neuroscience 38, 4482 (2018).

[54] L. A. Welikovitch, S. Do Carmo, Z. Maglóczky, P. Szocsics, J. Lőke, T. Freund, and A. C. Cuello, Acta neuropathologica 136, 901 (2018).

[55] G. W. Van Hoesen and B. T. Hyman, Progress in brain research 83, 445 (1990).

[56] G. W. Van Hoesen, B. T. Hyman, and A. R. Damasio, Hippocampus 1, 1 (1991).

[57] C. Pennanen, M. Kivipelto, S. Tuomainen, P. Hartikainen, T. Hänninen, M. P. Laakso, M. Hallikainen, M. Vanhanen, A. Nissinen, E.-L. Helkala, et al., Neurobiology of aging 25, 303 (2004).

[58] G. B. Chand, I. Hajjar, and D. Qiu, Human Brain Mapping 39, 4987 (2018).

[59] G. B. Chand and M. Dhamala, Neuroimage 134, 85 (2016).

[60] G. B. Chand and M. Dhamala, Neuroimage 152, 381 (2017).

[61] G. B. Chand, J. Wu, I. Hajjar, and D. Qiu, Brain connectivity 7, 401 (2017).

[62] G. B. Chand, J. Wu, D. Qiu, and I. Hajjar, Frontiers in aging neuroscience 9, 177 (2017).

[63] T. Timmers, R. Ossenkoppele, S. C. Verfaillie, C. W. van der Weijden, R. E. Slot, L. M. Wesselman, A. D. Windhorst, E. E. Wolters, M. Yaqub, N. D. Prins, et al., Neurobiology of Aging 79, 50 (2019).

[64] A. J. Aschenbrenner, B. A. Gordon, T. L. Benzinger, J. C. Morris, and J. J. Hassenstab, Neurology 91, e859 (2018).

[65] S. Gill, P. Mouches, S. Hu, D. Rajashekar, F. P. MacMaster, E. E. Smith, N. D. Forkert, Z. Ismail, A. D. N. Initiative, et al., Journal of Alzheimer’s Disease 75, 277 (2020).

[66] C. James, J. M. Ranson, R. Everson, and D. J. Llewellyn, JAMA network open 4, e2136553 (2021).

